# The developing kidney actively negotiates geometric packing conflicts to avoid defects

**DOI:** 10.1101/2021.11.29.470441

**Authors:** Louis S. Prahl, John M. Viola, Jiageng Liu, Alex J. Hughes

## Abstract

The physiological functions of several organs rely on branched tubular networks, but little is known about conflicts in development between building enough tubules for adequate function and geometric constraints imposed by organ size. We show that the mouse embryonic kidney epithelium negotiates a physical packing conflict between tubule tip duplication and limited area at the organ surface. Imaging, computational, and soft material modeling of tubule ‘families’ identifies six geometric packing phases, including two defective ones. Experiments in kidney explants show that a retrograde tension on tubule families is necessary and sufficient for them to avoid defects by switching to a vertical orientation that increases packing density. These results reveal developmental contingencies in response to physical limitations, and create a framework for classifying kidney defects.

**One-Sentence Summary:** Epithelial branching in the kidney causes a geometric packing conflict that is resolved through internally generated tensions

## Main Text

Epithelial organs perform diverse functions - lung alveoli exchange gases, kidney tubules exchange water and ions, and mammary acini secrete milk. However, little is known about how developmental constraints and compromises affect adult organ function per unit volume (*1*–*3*). In the embryonic mouse kidney, the branching ECAD^+^/calbindin^+^ ureteric epithelium creates an arborized network of epithelial tubules linked by junctions or ‘nodes’ (**Fig. 1A,B**) through reciprocal signaling interactions with SIX2^+^ cap mesenchyme and PDGFRA^+^ stroma (*4*). Ureteric tubule tips induce nephron formation (*5*) (**Fig. 1B**), so the eventual filtration and reabsorption function of the adult kidney likely scales with the number of tips. However, while some organs like the lung fill 3D space during branching, ureteric branching occurs at the kidney surface (*6*–*8*). This should create a conflict between the available organ surface area and exponentially increasing tip number (**Supplementary Text**). How then does the kidney navigate this conflict?

**Fig. 1.**
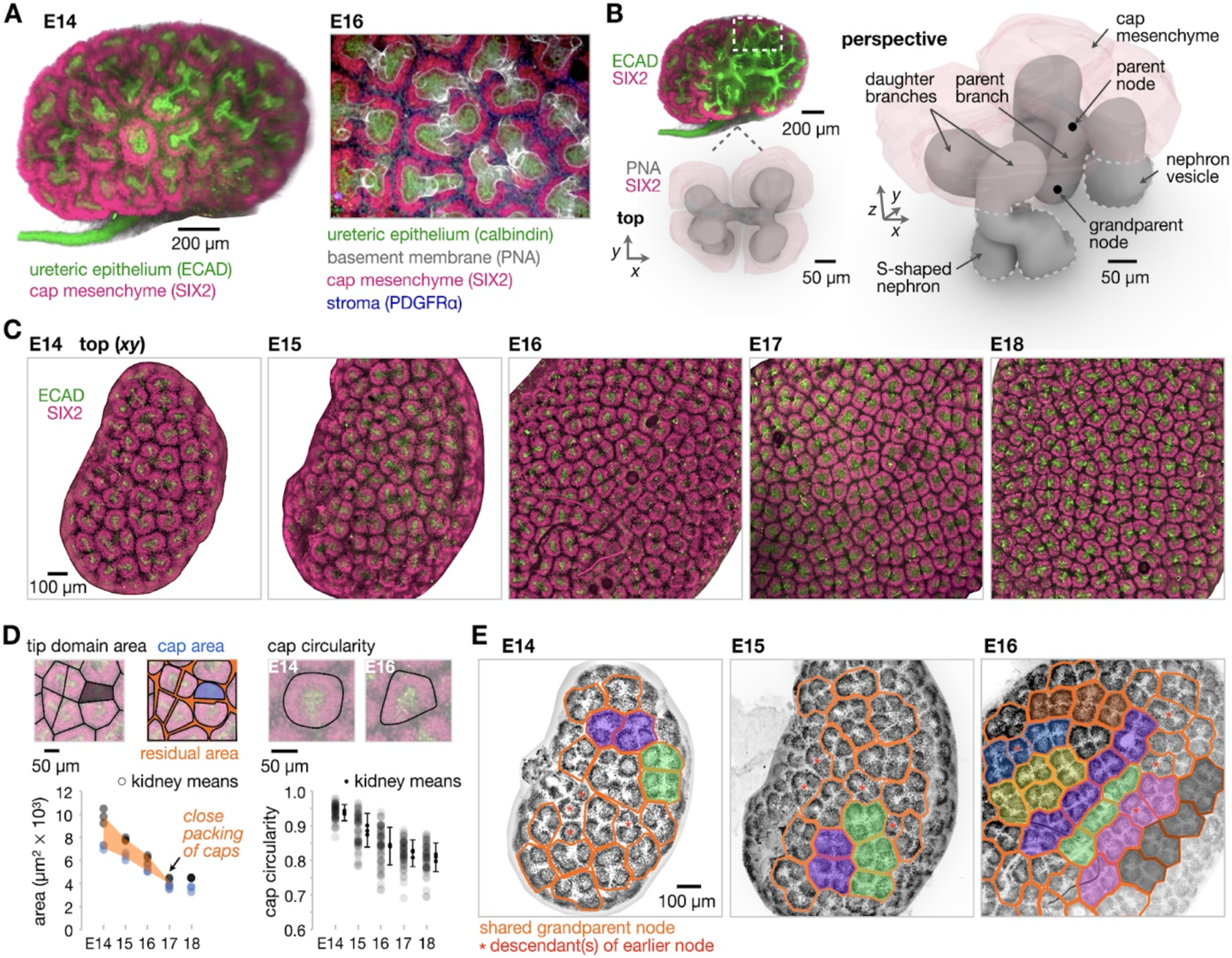
Tips of the ureteric epithelial tree form diverse patterns, pack closely, and achieve long-range order. (**A**) *Left*, 10x confocal projection of cleared mouse kidney at embryonic day (E)14. *Right*, detail of ureteric tips at E16. (**B**) Reconstruction of tubule family sharing a grandparent node. (**C**) 10x projections of kidneys at fixed zoom. (**D**) Quantitation of: *left*, tip domain (total area per tip) and cap mesenchyme area (means of >12 measurements per *n* = 3 kidneys per time point); *right*, cap mesenchyme circularity (>18 measurements per *n* = 3 kidneys per time point). Error bars are ± S.D. for all measurements per embryonic day. (**E**) Outlines of tubule families sharing a grandparent node. Fills indicate regions of aligned families.

To investigate tip patterning as branching progresses, we examined whole mount mouse kidneys at developmental stages from embryonic day (E)14-E18 by confocal immunofluorescence (**Fig. 1C**). Each ureteric tip is surrounded by a dynamic swarm of cap mesenchyme cells within distinct domains near the surface (*9*). We quantified the tip domain area as well as the SIX2^+^ cap area and circularity at each stage (**Fig. 1D**). Tip domain area and cap area both steadily decreased with organ stage, converging after E16. This indicates closer packing of cap mesenchyme populations due to a reduction in residual area occupied by stroma and parent nodes (junctions between two daughter tips), which appeared to drop into the kidney volume more quickly after E15.5. Cap circularity also steadily decreased since caps conformed to the area available to them as neighboring ones encroached. This indicates a compromise between mutual tip repulsion and confinement caused by crowding of neighboring tips over time (**Supplementary Text**).

In similar physical problems, repulsive objects can adopt distinctive packing geometries or ‘phases’, with critical transitions between them depending on the amount of confinement (*10*–*12*). We observed tubule families sharing a grandparent node progressively pack into aligned regions along the surface on the scale of 3-5 families at around E15-E16 (**Fig. 1E, Fig. S1**). We also observed dislocations and disclinations between aligned regions, which are hallmarks of liquid crystals and elastic crystalline solids (*13*, *14*). These data and published accounts of a lack of stereotypy (*7*, *15*) argue against a strict morphogenetic control program and instead for tip patterns being determined by competition between crowding and repulsion.

Can overcrowding of tips at the kidney surface cause geometric packing errors? If so, does the developing organ restructure tubule families to avoid these? To answer this, we made a computational model governed by three spatial parameters: 1) repulsion distance between tip centers, 2) planar confinement (width and depth) in the *xy* plane of tubule families each sharing a great-grandparent node, and 3) radial height occupied by families in *z* (**Fig. 2A, Movie S1, Supplementary Text**). The model uses energy minimization to predict branch positions at steady state (**Movie S2**), rather than simulating branching itself. A parameter sweep revealed six phases of tubule packing with distinct morphologies, plus one trivial case where tubules are inefficiently packed (**Fig. 2B, Fig. S2A**). Three phases are defined by parent nodes sitting near the kidney surface - ‘grandparent nodes at surface’, ‘parent nodes at surface’, and ‘H’s’. The H’s phase among these exhibits crystalline packing of families with long-range alignment. Two defective packing states appear as planar confinement increases. We call these states ‘short circuits’ where tubules intersect near the surface and ‘buried tips’ where surface overcrowding displaces some tips beneath the surface (**Fig. 2B, Supplementary Text**). Finally, a ‘vertical packing’ phase sees tips increasing their packing density by reorienting along the *z* axis (**Fig. S2B**). Published examples of tip patterns qualitatively reflect increased orientational ordering of tips in *xy* over ~E13.25-E16.5, and a transition to vertical orientation after ~E16, in step with corresponding model cases (**Fig. 2C**).

**Fig. 2.**
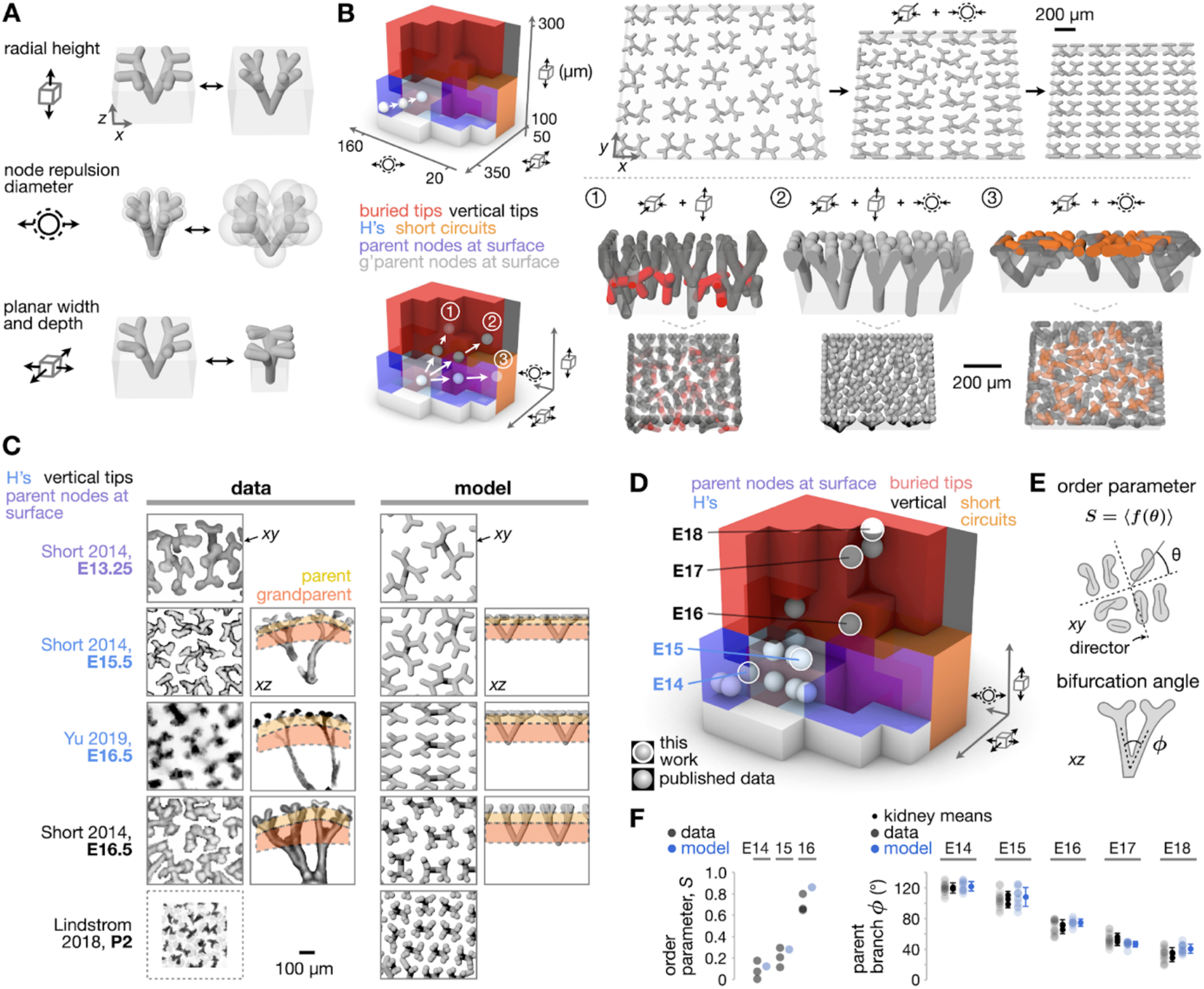
Published and measured kidney data map to a trajectory through physics-based model space that avoids defects. (**A**) Model output demonstrating geometric parameters. (**B**) *Left*, 3D plot of six phases of tip patterns at the model surface. *Top row*, an example trajectory from low tip density in the parent-nodes-at-surface phase into the H’s phase. *Bottom row*, three possible trajectories from the H’s phase into the buried tips, vertical, and short circuits phases. Insets show side views of tip patterns where red segments are buried in the model volume, and orange segments intersect with other segments. (**C**) *Left*, Top and side views, where available, of published ureteric tubule tip patterns. *Right*, corresponding views from model cases with matching geometric parameters. (**D**) Published kidneys and kidneys measured in this work (means of *n* = 3 kidneys per embryonic day) mapped to model phase space (see **Fig. S3** for full detail). (**E**) Quantitative metrics. (**F**) Quantitation of metrics comparing kidneys from this work and model cases. Error bars at right are ± S.D. for all measurements pooled across *n* = 3 kidneys for each embryonic day.

We quantified this by mapping spatial parameters from our own and published images of tubule families to the model, finding that kidney development roughly progresses along a trajectory in phase space through the H’s phase and ending with vertical packing at E17-E18 (**Fig. 2D, Fig. S3**) (*6*, *16*, *17*). We then evaluated two metrics of tubule organization. First, an order parameter *S* that measures local daughter branch alignment along orthogonal preferred directions (directors) when parent nodes are visible near the surface (**Fig. 2E**). *S* = 1 indicates crystalline packing and perfect ordering of each daughter along either director (**Supplementary Text**). *S* increased from E14-E16 as tubules entered the H’s phase with close agreement between experiment and model (**Fig. 2F**). Second, we quantified the parent branch bifurcation angle (*ϕ*) (**Fig. 2E**) and found that this progressively decreased in both experiment and model between E14 and E18 as tips shifted toward vertical packing (**Fig. 2F**) (*16*, *18*). The packing model thus explains developmental variation in tip patterns. It also predicts that a vertical packing switch is necessary for the developing kidney to increase tip density while avoiding defects.

We next asked whether physical forces could mediate the switch to vertical packing, since this requires that parent through great-grandparent nodes move further from the kidney surface (note increase in yellow and orange layer heights in **Fig. 2C**). Lindström *et al*. described ‘node retraction’ as retrograde movement of branch points towards the ureter beginning at ~E15 (*19*), which is consistent with our data showing a reduction in *ϕ* during the shift to vertical packing (**Fig. 2D,F**). However, it is unclear whether this rearrangement is caused by revision of branch point locations through collective cell dynamics or by an active retraction force generated within the ureteric epithelium. We reasoned that a retrograde force on tubules (Fz) could be indirectly measured by disrupting the basement membrane at the epithelial-mesenchymal interface using dispase (*20*). This would release adhesive and shear forces at that interface and cause tips to rapidly retract away from the kidney surface (**Fig. 3A**). Indeed, dispase added to E15 kidney explants caused basement membrane disruption, epithelial delamination from the surrounding mesenchyme and tip and parent node retraction away from the kidney surface in <10 mins (**Fig. 3B,C, Movie S3**). We found significant decreases in the tip areas at the surface relative to untreated controls that were sensitive to either the myosin II inhibitor blebbistatin or the ROCK inhibitor Y-27632 (**Fig. 3C**). In contrast, E14 kidneys showed less tubule retraction upon dispase treatment, but returned to the retraction level of E15 kidneys in the presence of calyculin A (calA), which stimulates actomyosin contractility (*21*) (**Fig. 3C**). These data indicate that the epithelial-mesenchymal interface is subject to an actomyosin-based retraction force that switches on at ~E15 during the shift to vertical packing.

**Fig. 3.**
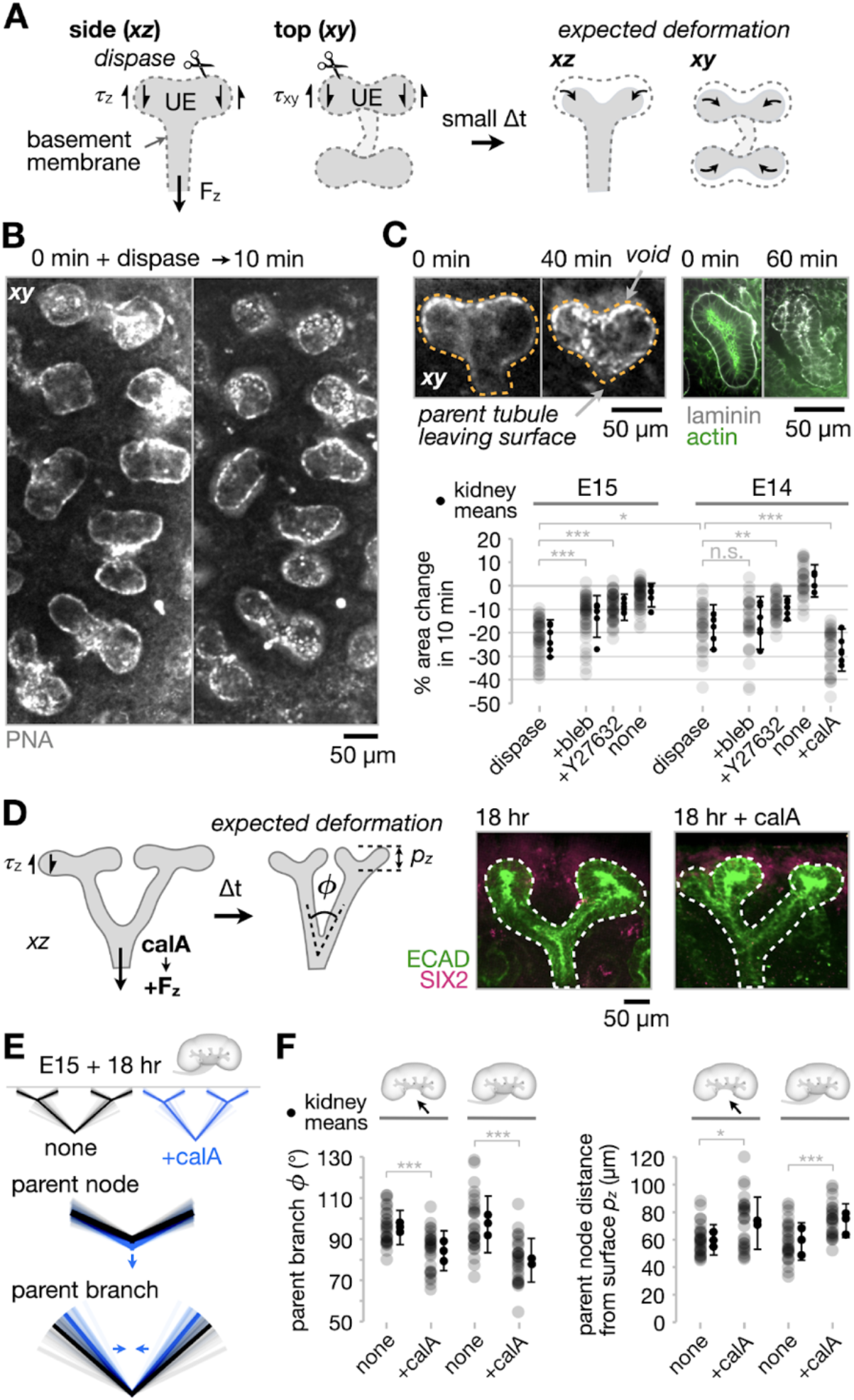
Retrograde tubule tension is sufficient for transition to vertical packing. (**A**) Expected tip deformation after dispase. (**B**) Tip pattern in live E15 kidney explant before (*left*), and after 10 min dispase (*right*). (**C**) *Top left*, Example daughter tip pair. *Top right*, Laminin (basement membrane) immunofluorescence for similar pairs +/− dispase. *Bottom*, Tip area change after 10 min dispase; 60 min pre-treatment with 20 μM blebbistatin (bleb), 20 μM Y-27632, or 25 nM calyculin A (calA) followed by 10 min dispase; or no dispase (none) for E14 and E15 kidneys (>5 measurements per *n* = 6 kidneys per condition). Error bars are ±S.D. across all measurements per condition. (**D**) *Left*, Expected tubule deformation with calA; *ϕ* = parent bifurcation angle, *p*_*z*_ = parent node distance from kidney surface. *Right*, Control and calA-treated tubule families after 18 hr culture. (**E**) Pictographs of family deformation using *ϕ* and *p*_*z*_ data from (F). Transparent lines are individual measurements, solid lines are means. (**F**) Quantitation of *ϕ* and *p*_*z*_ for kidneys +/− ureter, and +/− calA for 18 hr (5 measurements per *n* = 3 kidneys per condition). Error bars are ±S.D. across all measurements per condition. (C),(F) One-way ANOVA, Tukey’s test (\**p* < 0.05, \*\**p* < 0.01, \*\*\**p* < 0.001).

Is this retraction force then sufficient to shift tubule families from the crystalline H’s state to more vertical packing? To test this, we added calA to intact E15 kidneys and cultured for 18 hr. We then measured *ϕ* and the radial distance of parent nodes from the kidney surface (*p*_*z*_) as metrics for the vertical transition of tip families (**Fig. 3D**). Indeed, *ϕ* decreased and *p*_*z*_ increased, whether or not the ureter had been removed (**Fig. 3E,F**). This revealed that forces intrinsic to the ureteric tree are sufficient to explain the transition of tubule families to the vertical packing phase.

If geometric crowding, tip repulsion, and tubule tension are primarily responsible for tip positions, similar outcomes should translate to material systems with similar geometry. We therefore established a biomimetic soft material model of ~E16 ureteric epithelial families that each share a great-great-grandparent node using 3D printed elastic filaments embedded in a soft silicone elastomer (**Fig. 4A, Movie S4**). Applying compressive strains in *xy* that mimic tip crowding by duplication and elongation strains in *z* were sufficient to create overlapping tip defects and vertical tip transitions, even in a purely abiotic system (**Fig. 4B-D, Supplementary Text**).

**Fig. 4.**
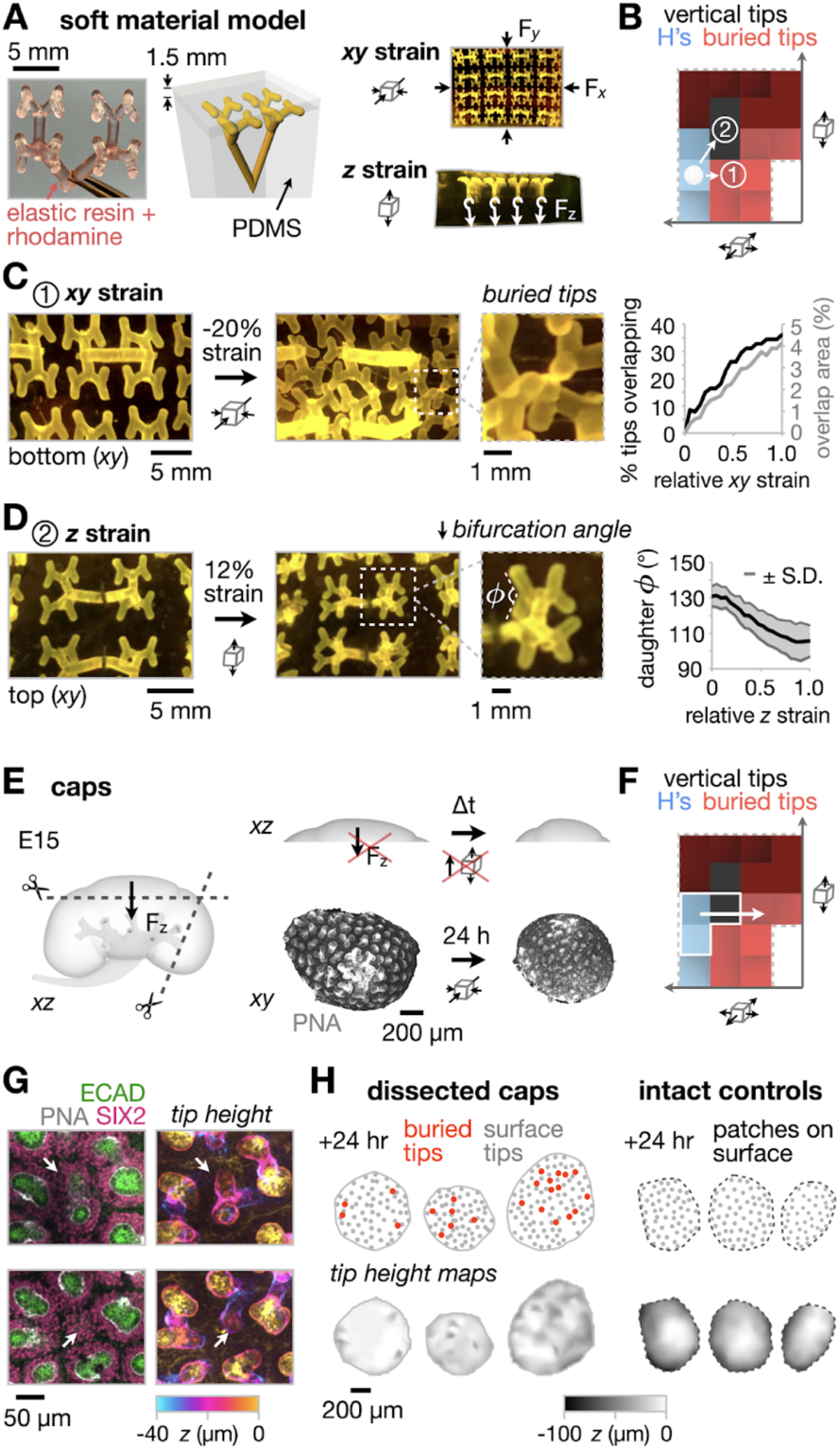
Retrograde tubule tension is necessary to avoid buried tips. (**A**) *Left*, 3D-printed elastic tubule family. *Middle*, Rendering of embedding in polydimethylsiloxane (PDMS) silicone. *Right*, Top and side-view of tubule family array and deformation modes. (**B**) 2D section of phase space showing expected transitions under strain in *xy* and *z*. (**C**) *Left*, Photographs of tubule positions taken from underside of the model before and after strain is applied in *xy*. *Right*, Quantitation of tip overlap metrics. (**D**) *Left*, Similar photographs for added strain in *z*. *Right*, Quantitation of daughter branch bifurcation angle with ± S.D. envelope. (**E**) *Left and top*, cartoon of cap dissection and change in strain parameters over time in culture. *Bottom*, confocal projections of a cap before and after 24 h culture. (**F**) 2D section of phase space showing expected transition for caps in culture. (**G**) Buried tips shown as *left*, confocal immunofluorescence image, *right*, depth-coded projection; one example per image row. (**H**) *Top*, pictographs of surface and buried tip locations after 24 h culture. *Bottom*, tip height maps interpolated from (*x,y,z*) positions of tips.

We then attempted to force a forbidden phase transition predicted by the models in live explants by removing F_z_ from the ureteric tree during the transition to vertical tubule orientation. We dissected ‘caps’ of cortex from E15 kidneys to sever tubules (and remove force transmission) roughly below the great-grandparent nodes and cultured them for 24 hr (**Fig. 4E**). These caps compacted in area in the *xy* plane by 10 ± 7% (*n* = 6 caps, **Table S1**), accentuating planar confinement of tubule families. Our computational and soft material models predict that this would create buried rather than vertical tips near the kidney surface (**Fig. 4F**). Accordingly, we found that 34 of 290 tips were buried below the surface of dissected caps after 24 hr (across *n* = 4 caps), whereas none of 161 tips were buried in intact control kidneys (across *n* = 4 kidneys) (**Fig. 4G,H**). Buried tips were visible as ‘dimples’ in tip height maps for dissected caps after 24 hr, relative to smooth maps for intact controls and caps fixed immediately after cutting (**Fig. 4H, Fig. S4**). This experiment revealed that retraction forces on tubules are necessary for the ureteric tree to resolve into vertical packing without generating buried tip defects.

Finally, we wondered if our model could predict the impacts of developmental defects on the geometry of the ureteric tree. Genetic mutations and environmental factors including prematurity and nutrient deficiency can affect kidney size and nephron number (*3*, *22*, *23*), branch organization of the ureteric tree (*24*), and the risk of fetally programmed adult diseases (*3*). We successfully mapped three published mutants to regressive or forbidden model phases (**Fig. S5, Supplementary Text**), revealing that the model morphospace may enable insight into abnormal cell and tissue dynamics contributing to an observed tree morphology.

Here we use physics-based simulations, soft material modeling, and kidney explants to reveal a conflict between kidney surface area and exponential duplication of ureteric epithelial tips there. One outcome of this conflict is the emergence of defective packing states. These must be actively avoided through tubule tension toward vertical tip packing. We propose that similar conflicts between increased function and geometric constraints may be at play in other complex organ systems, perhaps amplifying the structural consequences of congenital defects.

## General

We would like to thank Hughes lab members and instructors and students of the 2021 MBL Embryology course for training, discussion and advice. We are grateful to A. Gupta, K. Sasaki, and Y. Sakata for technical assistance with kidney dissections; and to L. Bugaj, A. Raj, and W. Marshall for feedback on the manuscript.

## Funding

This work was supported by:

NIH F32 fellowship DK126385 (LSP)

Predoctoral Training Program in Developmental Biology T32HD083185 (JMV)

NIH MIRA R35GM133380 (AJH)

NSF CAREER award 2047271 (AJH)

## Author contributions

Conceptualization: LSP, AJH

Methodology: LSP, JMV, JL, AJH

Software: AJH

Formal analysis: LSP, AJH

Investigation: LSP, JMV, JL, AJH

Writing - original draft: LSP, AJH

Writing - review and editing: LSP, JMV, JL, AJH

Visualization: LSP, AJH

Supervision: LSP, AJH

Project administration: AJH

## Data and materials availability

All data necessary to evaluate conclusions of this study are presented in the paper and supplementary materials. In-house code and raw data are available from the authors upon request.

## Supplementary Materials

Materials and Methods

Supplementary Text

Figs. S1 to S5

Table S1 Movie S1 to S4

Computational model file

