## Supplementary Materials for "The developing kidney actively negotiates geometric packing conflicts to avoid defects"

**This PDF file includes:**

Materials and Methods
Supplementary Text
Figs. S1 to S5
Table S1
Captions for Movies S1 to S4
Caption for Data S1

**Other Supplementary Materials for this manuscript include the following:**

Movies S1 to S4
Data S1 (computational model file)

### Materials and Methods

Animal experiments

All mouse experiments followed NIH guidelines and were approved by the Institutional Animal Care and Use Committee of the University of Pennsylvania. Embryos were collected from timed pregnant CD-1 mice (Charles River) at developmental stages between embryonic day (E)14 and E18, dissected in chilled Dulbecco's phosphate buffered saline (DPBS, MT21-31-CV, Corning), and kidneys were fixed with ice cold 4% paraformaldehyde (PFA, 16% stock, 15710, Electron Microscopy Sciences) diluted in DPBS for 12-20 min or further processed for live analysis. Developmental kidney stages were roughly confirmed by limb staging of whole embryos, as previously described (*16*, *25*). In some experiments, whole embryos were fixed overnight in 4% PFA at 4°C prior to dissection. For cap experiments, E15 kidneys were dissected using a razor blade to retrieve lateral border and superior pole tissue segments. Kidneys and caps were labeled with 20 µg $ml^{-1}$ AlexaFluor 488-labeled peanut (*Arachis hypogaea)* agglutinin lectin (PNA, L21409, Sigma) in Dulbecco's minimum essential medium (DMEM, 10-013-CV, Corning) for 60 min, washed in 2 exchanges of DMEM and cultured for 24 hr at 37°C, 5% $CO_2$ in microplate wells coated with 1% agarose prior to fixing, immunofluorescence, and confocal imaging. For dispase experiments, whole E14-E15 kidneys were PNA lectin labeled with or without drugs for 60 min, washed with media and transferred to 2 mm-diameter wells to reduce sample movement during dispase addition. These wells were created with a biopsy punch in a ~5 mm-thick layer of 15:1 (base:crosslinker) polydimethylsiloxane (PDMS) elastomer (Sylgard 184, 2065622, Ellsworth Adhesives) set in 35 mm coverslip-bottom dishes (FD35-100, World Precision Instruments). A 1:1 mix of 50 U $ml^{-1}$ dispase (Corning 354235):DMEM was added immediately prior to time-lapse imaging. Dispase experiments included co-treatment with myosin phosphatase inhibitor calyculin A (25 nM, 9902S, Cell Signaling Technology), myosin II ATPase inhibitor (S)-(-)-blebbistatin (20 µM, 1852, Tocris Bioscience), or RHO/ROCK inhibitor Y-27632 hydrochloride (20 µM, 72304, STEMCELL

Technologies). Calyculin A and blebbistatin were diluted in dimethyl sulfoxide (DMSO, D2650, Sigma) and stored in working aliquots (10 µM and 30 mM, respectively) at -20°C and diluted in PNA lectin labeling media immediately prior to use. Y-27632 was diluted in sterile DPBS at 10 mM, stored in working aliquots at -20°C, and similarly diluted in labeling media. PNA lectin was diluted to a 5 µg $ml^{-1}$ stock in DI water and stored in working aliquots at -20°C.

Tip packing models

Tip packing simulations were performed using Kangaroo2 (Daniel Piker), a position-based dynamics solver within the Rhino Grasshopper algorithmic modeling environment (Robert McNeel & Associates). We took a form-finding approach here to enable rapid local solution to the packing problem for a 6 x 4 array of tubule families each sharing a great-grandparent node confined to a rectangular cuboid. The Kangaroo2 solver considers forces at model vertices caused by geometric goals and moves points to satisfy force balances through dynamic relaxation (*26*, *27*), reducing the sum of energy potentials associated with each goal while employing momentum and damping to allow models to transiently explore local solutions during convergence. Tubule families were constructed from nodes and edges at a scale of 85 µm per model unit. Daughter, parent, and grandparent tubule lengths show little temporal variation across E13.5-E16.5 and were set at 57, 73, and 120 µm respectively for all model cases to roughly match those in the mouse kidney (*16*). Model parameters were created in Kangaroo2 with relative energy potential weightings called 'strengths'. All edges were modeled as linear elastic elements with an arbitrarily high strength of $K$ = 100 and rest lengths equal to their initial lengths. Nodes were repulsive with strength $R$ = 0.8 and diameter $D$. Nodes received a vertical load toward the model kidney surface of strength $V$ = 0.5. Strengths controlling node repulsion from the cuboid boundaries and anchoring great-grandparent nodes to the lower boundary were both set arbitrarily high at 100 and 1000 respectively. $R$ and $V$ parameters are not readily measurable from kidney samples, instead they were manually adjusted to qualitatively fit the models to tubule families' bifurcation angles in E16.5 kidney data from Yu *et al.* and then fixed for all other simulations (*6*). The model phase diagram was constructed by sweeping through an 'input space' (*xy* extent of the confining cuboid, *z* extent of the cuboid, and repulsion diameter $D$) and measuring and plotting the 'output space' phase diagram as a function of 1) the square root of the area of tip families at the cortical surface (~planar width and depth of confinement), 2) the radial height of tip families, and 3) the apparent repulsion distance between tip centers. Phases were assigned by qualitative assessment of tip patterns in each of the 216 simulations. All other tip families from experiments and literature were mapped to the model output space by measuring the same three output parameters (averaged across at least 3 tip families per kidney) and finding the nearest simulation case having the smallest sum of squared error across the three parameters.

Soft material model

Tubule families sharing great-great-grandparent nodes were modeled in Rhino to approximately match their dimensions in kidney trees from Yu *et al.* at E16.5 (*6*). Models were scaled up isotropically by a factor of 30 to meet minimum resolution requirements for subsequent 3D printing in elastic resin (50A Shore durometer hardness, 3.23 MPa ultimate tensile strength) using a Form 3 printer (FormLabs). Tubule models were impregnated with 10 µM methacryloxyethyl thiocarbamoyl rhodamine B (Polysciences 23591) in acetone for 10 min, washed with DI water and dried for 1 hr at room temperature. The models were then placed in a 4 x 4 array on a 1.5 mm layer of 50:1 (pre-polymer base:curing agent) PDMS degassed under vacuum and cured in a 8 x 8 x 3.5 cm plastic container at 60°C for 2 hours. A second layer of degassed 50:1 PDMS was then poured until great-great-grandparent nodes were just exposed above the PDMS surface, and then cured at 60°C overnight. 1# (4 x 7.5 mm) fishing hooks pre-tied with nylon monofilament line were secured to each great-great-grandparent node, and a final 50:1 PDMS layer was poured for a total model height of 3 cm. Nylon lines were held vertically over great-great-grandparent nodes during curing at 60°C overnight before demoulding the completed model from the plastic container.

Immunofluorescence and optical clearing

Whole-mount immunofluorescence staining of whole kidneys and cap segments was adapted from Combes *et al.* and O'Brien *et al.* (*28*, *29*). Briefly, fixed kidneys were washed 3 x 5 min in ice cold DPBS, blocked for 2 hr at room temperature in PBSTX (DPBS + 0.1% Triton X-100) containing 5% donkey serum (D9663, Sigma), incubated in primary and then secondary antibodies in blocking buffer for at least 48 hr at 4°C, alternating with 3 washes in PBSTX totaling 12-24 hours. Minimum duration of primary and secondary incubations and washes depended on the age of the kidney, as described (*28*). The E14 example kidney in **Fig. 1A-B** and all kidneys used to measure parent branch angles and node distances from the kidney surface in **Figs. 2-3** were cleared for 2 days in ScaleA2 (4M urea + 0.1% Triton X-100 + 10% glycerol), followed by 2 days in ScaleB4 (8M urea + 0.1% Triton X-100) (*30*) and were imaged in 35 mm culture dishes or 24-well glass bottom plates containing ScaleA2. Tissue clearing by ScaleA2/B4 caused a 33 ± 6% increase in overall kidney cross sectional area measured by brightfield or fluorescence microscopy, similar to previously reported values of tissue expansion from this clearing method (*30*).

Primary antibodies and dilutions included rabbit anti-Six2 (1:600, 11562-1-AP, Proteintech, RRID: AB_2189084), mouse anti-E-cadherin clone 34 (1:200, 610404, BD Biosciences, RRID: AB_397787), mouse anti-calbindin D-28K clone CB-955 (1:100, C9849, Sigma, RRID: AB_476894) rat anti-CD140/PDGFRA (1:200, 14-1401-82, eBioscience, RRID: AB_467491), and rabbit anti-laminin (1:50, ab11575, Abcam, RRID: 298179). Secondary antibodies (all raised in donkey) include anti-rabbit AlexaFluor 647 (1:300, A-31573, ThermoFisher, RRID: AB_2536183), anti-rabbit AlexaFluor 555 (1:300, A-31570, ThermoFisher, RRID: AB_2536180), anti-mouse AlexaFluor 555 (1:300, A-31572, ThermoFisher, RRID: AB_162543), and anti-rat AlexaFluor 405 (1:300, A48268, ThermoFisher). In some experiments, samples were counterstained in 300 nM DAPI (4',6-diamidino-2-phenylindole; D1306, ThermoFisher), 20 µg $ml^{-1}$ PNA lectin-Alexa 488, and/or 1:40 AlexaFluor 647 phalloidin (A22287, ThermoFisher) diluted in blocking buffer for 2 hours at 4C prior to imaging.

Imaging and data analysis

Kidney imaging was performed using a Nikon Ti2-E microscope equipped with a CSU-W1 spinning disk (Yokogawa), a white light LED, laser illumination (100 mW 405, 488, and 561 nm lasers and a 75 mW 640 nm laser), a Prime 95B back-illuminated sCMOS camera (Photometrics), motorized stage, 4x/0.2 NA, 10x/0.25 NA and 20x/0.5 NA lenses (Nikon), and a stagetop environmental enclosure (OkoLabs). In dispase treatment experiments, a 150 µm stack (21 frames, 7.5 µm step size) of the kidney cortex was collected at each stage position every 5 minutes. Z-stacks of fixed and stained dispase-treated kidneys or cut caps (and controls) were similarly collected using 5 µm step size (100-200 µm, 21-41 frames) using either the 10x or 20x lens. In order to image the cortical kidney surface, caps were oriented cortex-side against the glass on 35 mm dishes while uncut controls were oriented cortex-side down in the bottom of 2 mm-diameter PDMS wells, as described above. The soft material model was imaged by submerging it in a transparent glass water bath and filming through an amber trans-illuminator filter (IO Rodeo) during manual manipulations under blue 470 nm LED illumination (ThorLabs M470L2-C1) using an iPhone 8 video camera.

Tubule tips at the kidney surface in dispase experiments were outlined by manually tracing the outline of PNA lectin signal using the *polygon sections* tool in FIJI. Percentage area change was defined using the difference in recorded areas between the $t$ = 0 and $t$ = 10 min timepoints. We used a similar procedure to quantify kidney or cap area change due to cutting, calA treatment, ureter removal, or swelling from exposure to tissue clearing solvents. Tip heights in whole kidneys and caps were manually annotated in FIJI by marking ($x$,$y$,$z$) coordinates at the center of tip lumens using the *multi-point* tool. Coordinates were translated into height maps using cubic interpolation via in-house MATLAB code. Soft material model strains, tip overlap, and tip angles were manually measured in FIJI. Parent branch bifurcation angles and node distances from the kidney surface were manually annotated from xz sections of cleared kidney data and model renderings.

For orientation analysis, we made use of order parameter metrics previously applied to liquid crystals (*31–33*):

For 2D: $S = \left\langle 2\cos^2(\theta - \theta_d) - 1 \right\rangle$

Where $\theta$ is the orientation of an object and $\theta_d$ is the orientation of the 'nematic director', or the apparent preferred orientation of objects in a local field of view that serves as a reference axis. However, this metric cannot be directly applied. Since bifurcations are asynchronous, some new daughter branches are approximately orthogonal to nearby branches that have not yet bifurcated. This implies that the degree of order in daughter orientations should relate to their alignment along either orthogonal director. Indeed, histograms of daughter branch angles in the *xy* plane manually annotated in FIJI showed their overall orientations to be bimodal, with modes ~$90^o$ apart, such that directors for order parameter quantitation could be defined as the center of each mode. We therefore modified the 2D expression for $S$ by adding a factor of 2 to the argument of $\cos^2(\theta)$, allowing $S$ to quantify alignment along two orthogonal directors. Our orientational order parameter for daughter branches was therefore:

$$S = \left\langle 2\cos^2(2(\theta - \theta_{\mathrm{Mo}})) - 1 \right\rangle$$

Where $\theta$ are daughter branch angles and $\theta_{\mathrm{Mo}}$ is the orientation of one of the directors ($\theta,\theta_{\mathrm{Mo}} \in$ [0,180]). This order parameter will therefore be $S$ = 0 for random orientation of daughters, and $S$ = 1 for perfect alignment of each daughter along either director.

### Supplementary Text

Surface area limitation to ureteric branching and compromise between tip repulsion and confinement

Unlike the mammalian lung, ureteric tubule tips remain at the kidney surface throughout development (*6*, *8*). In the mouse, the number of tubule tips grows exponentially through tip doubling from the onset of bifurcations at ~E12.5 until branching stops at around postnatal day P2 (*16*, *34*). Here, we consider that the overall kidney surface area limits the total number of branching tips of finite size that can possibly be present at the surface, which can be addressed using a simple geometric argument. First consider the kidney as a growing sphere of radius $r$ with functional capacity proportional to its volume. The extra kidney surface area available to new tips then decreases proportional to ~$1/r$ per unit function, while the number of tips grows proportional to $r^2$ (*16*). This constraint sets up a conflict between the surface area availability and the number of tips of finite size that can possibly be present at the surface.

Other evidence comes from our observation of a marked decrease in the circularity of cap mesenchyme domains over E14-E18, which is caused by caps conforming to the area available to them as neighboring caps encroach. This indicates a compromise between tip repulsion and confinement caused by crowding together of neighboring tips over time. This compromise is also indirectly supported in other ways. Short *et al.* noted that the branch angle between daughter tubules measured close to their bifurcation point was higher than the angle measured relative to their tips after ~E15 (*16*), adding evidence that tips are shunted closer together due to crowding by neighbors. Davies *et al.* noted self-avoidance of ureteric tubules, since when kidney explants are cultured at the air-media interface they flatten due to gravity. This flattening would locally increase tubule tip confinement in some areas, but the ureteric epithelium still branches primarily in *xy* without tubules colliding (*35*). The same study implicated TGFβ family members (in particular BMP7) as contributing a portion of the tip repulsion effect in explant cultures, although self-avoidance defects were not observed in *Bmp7*$^{-/-}$ kidneys (*15*). Repulsion could also be a physical phenomenon caused by cell-cell adhesion, such that cap mesenchyme cell populations form an elastic barrier that resists infiltration of a nearby tip. Indeed, the nephron progenitors that make up each cap are thought to adhere more tightly to the ureteric epithelium and to each other than to the surrounding stroma (*36*, *37*).

We found that tip repulsion is crucial to prevent tubule collisions, but can also force tips off the surface and into the kidney volume. Our model assumes that repulsion between tubule tips is adequately captured by a mechanical analogy to compressing a Hookean spring. In reality, the cap mesenchyme is in a dynamic equilibrium, with significant cell motility within cap populations and between caps (*9*). These caps are also separated by sheets of stroma whose role in repulsion have not been systematically examined. Modeling repulsion as an elastic interaction is not intended to capture the biological details. Further study of mesenchymal dynamics could clarify which biochemical and potentially biophysical inputs instruct maintenance of a cohesive mesenchymal barrier between tips, and what processes enable changes in repulsion distance such as by thinning of the stroma. Sufficient control over tip repulsion would also allow us to test transitions to the short circuits phase more rigorously. Even so, a fixed elastic repulsion constraint appears to adequately predict tip geometry over a large range in developmental time and for the small set of published mutants that we examined.

Once ureteric tubules are vertically packed, our model suggests that there are no other structural remodeling options for increasing tip surface density without crossing into forbidden phases. If mouse kidneys are optimally packed, it would appear impossible for larger kidneys with more restrictive surface area:volume ratios to evolve. However, additional levels of hierarchical organization can overcome the geometric constraints imposed by packing nephrons into a single lobe (renicule) as in the mouse kidney. Indeed, human and bovine kidneys are composed of multiple lobes that all still share a border with the kidney surface, while lying above a proportionally more extensive system of papillae and calyces draining into the pelvis (*17*, *38*). This would increase the surface area:functional volume of the kidney to a point, at the expense of larger and larger drainage networks. Whales exhibit yet another shift in hierarchy - their kidneys contain close-packed renicules, each encapsulated and roughly the size of mouse kidneys, such that not all renicules occupy organ surface area, yet all individually maintain cortico-medullary structure (*38*, *39*). Thus, physical constraints appear to have exerted selective pressure on the morphology of tubule families and whole kidneys as the size of mammals increased across evolutionary time.

<u>Soft material model dynamics</u>

We 3D printed tubule families that share a great-great-grandparent node as soft elastic filaments, impregnated them with fluorescent rhodamine, and embedded them in arrays near the surface of a block of soft 50:1 (pre-polymer base:curing agent) PDMS silicone elastomer (**Fig. 4A, Movie S4**). The 3D geometry of these filament families and their spacing in *xy* was designed to mimic those of the ~E16 kidney in the H's phase. Since the average distance between ureteric tubule tips decreases by <30% from E15.5 to postnatal day 2 (*17*), the PDMS embedding medium mimics an approximately fixed repulsion distance between tips maintained by the mesenchyme and stroma, while the mechanical flexibility in each polymer allows tips to move in space and adopt energetically favorable positions and orientations. We next applied compressive strains in *xy* to simulate increasing competition of tips for space at the kidney surface as they bifurcate, and tension strains in *z* to simulate tubule retraction. The model predicts that these deformations would result in formation of buried and vertical tip outcomes respectively (**Fig. 4B**). Indeed, applying a -20% strain in xy to the silicone block led to rotation and deflection of some tips downwards, where they became buried under others (**Fig. 4C**). Applying a 12% strain on the filament families by pulling downwards on the great-great-grandparent nodes using nylon monofilaments lead to closer clustering of daughter branches and reduction in the bifurcation angle between sisters (**Fig. 4D**). These results show that physical interactions among tips and their embedding medium in the presence of strains acting at long range are sufficient to explain buried vs. vertical tip patterning transitions.

<u>Mapping kidney mutants to the model</u>

We mapped three mutants onto the model phases, as a first step toward an overarching geometric classification scheme. To do this, we measured the three spatial parameters in our model for three published perturbations to predict their effect on ureteric morphology (**Fig. S5A**). Indeed, Cer1 overexpression, diphtheria toxin (DTA)-induced partial ablation of the cap mesenchyme, and Spry1 knockout kidneys could all be mapped to model phases. Overexpression of the BMP antagonist Cer1 is known to induce ectopic 'side branches' in the ureteric epithelium, which were also reported in kidney explants after air-liquid interface culture (*40*). Indeed, although such culture conditions can distort kidney dimensions and branch morphology (*9*), Cer+ kidneys did fall into the buried tips phase at E11.5 + 4 days in culture as compared to the vertical packing phase for the wild-type cultured under the same conditions (**Fig. S5B**). Kidneys engineered to produce DTA in *Gdnf*-expressing cap mesenchyme cells become hypoplastic, despite retaining normal gross organization (*41*). The wild-type kidney presented in Cebrian *et al.* appeared to have the size, ureteric tip number, and daughter tubule orientational order parameter (*S*) properties of an ~E15-16 kidney, comparing against our own data and other publications. Even so, the corresponding E14.5 DTA kidney mapped to the parent-nodes-at-surface phase rather than the H's phase as for the wild-type, and showed a corresponding drop in *S* that was mirrored by model cases (**Fig. S5C**). This revealed that rather than DTA mutants simply having fewer tips in the same H's pattern as the wild-type, the mutant instead has fewer tips in the more amorphous packing pattern predicted for an *earlier* developmental stage. Finally, the GDNF/RET antagonist Sprouty (Spry1) knockout or knockdown kidneys tend toward poorly spaced tubules that occasionally appear cystic, incompletely branched, or fused, with more rapid branching and shorter tubule lengths between branch points (*15*, *42*–*44*). E13.5 Spry1$^{-/-}$ mutants mapped to the short circuits phase of the model rather than the H's phase as for the wild-type (**Fig. S5D**). This semi-quantitative agreement between model and published mutants suggests that geometric features may be used to map a range of mutants into a shared morphospace that would enable more targeted hypothesis testing regarding abnormal cell and tissue dynamics contributing to an observed tree morphology.

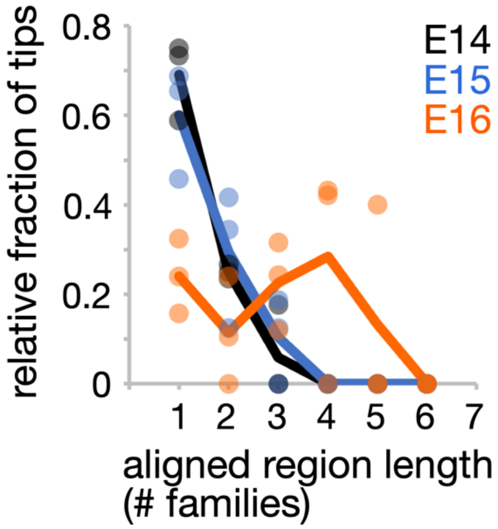


**Fig. S1. Long-range alignment of tubule families approaching the H's phase.** Quantitation of the relative fraction of tips associated with aligned regions of length listed on the *x* axis for E14-E16 kidneys (*n* = 3 kidneys per time point). Solid lines connect averages.

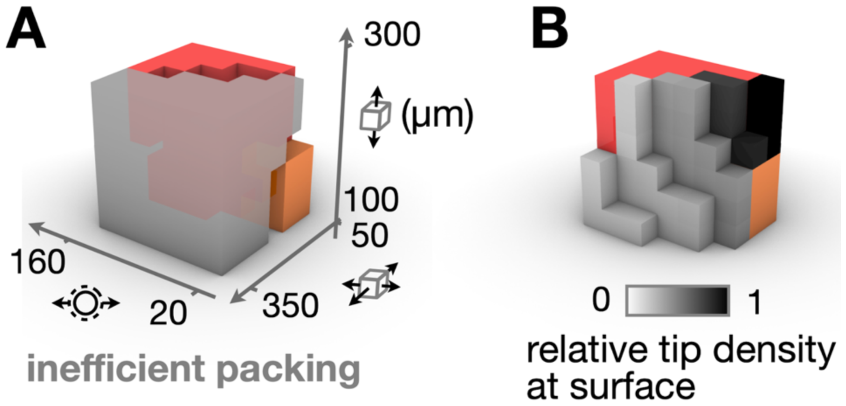


**Fig. S2. Tip packing phase diagram reveals inefficient packing phase and tip density advantage in the vertical phase.** *Left*, a trivial phase of inefficient packing where gaps on the order of a tubule family surface area appear. *Right*, relative density of tubule tips at the model surface in ‘allowable’ phases.

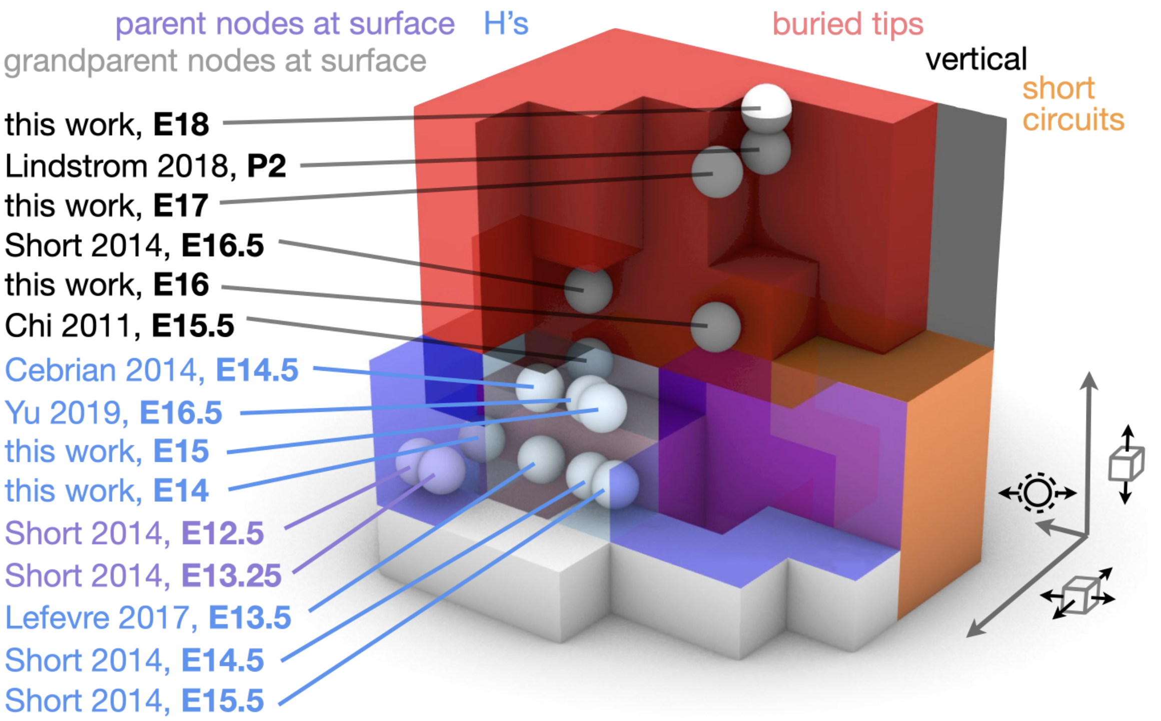


**Fig. S3. Full version of Fig. 2D.**

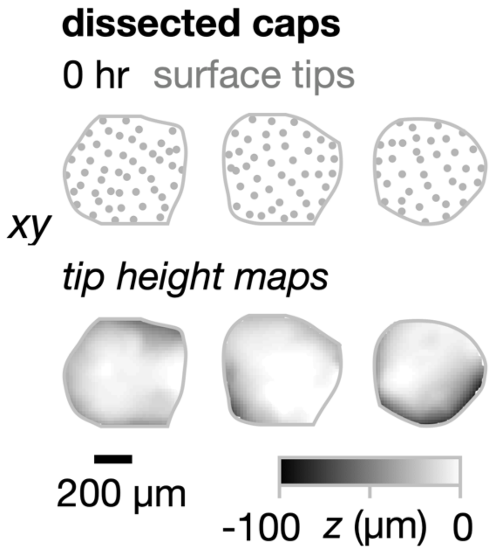


**Fig. S4. Cap cutting does not itself induce buried tips.** Pictographs of tip positions and tip height maps for E15 caps immediately after dissection.

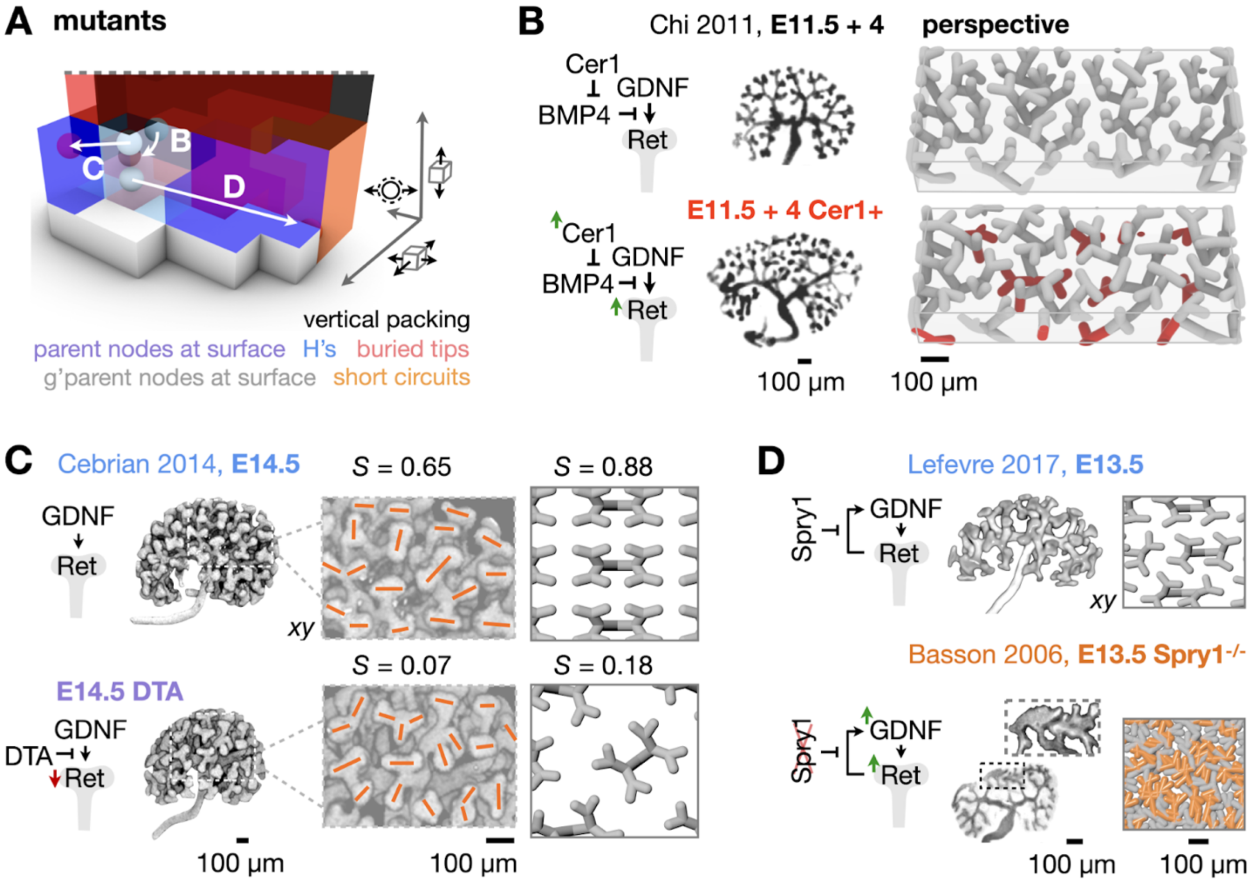


**Fig. S5. Published mutants map to regressive or forbidden model phases.** (**A**) A portion of the model phase diagram showing predicted packing phases based on geometric properties of normal control kidneys (white markers) compared to mutant kidneys in (B)-(D) (red markers). (**B**) *Left*, cartoons of Cer1 interaction with the GDNF/RET pathway central to branching morphogenesis. Middle, fluorescence micrographs reproduced from Chi *et al*. for E11.5 kidneys cultured for 4 days. *Right*, model predictions of branch geometry in wild-type and Cer1+ mutant. (**C**) Similar data for wild-type and diphtheria toxin (DTA)-induced cap mesenchyme ablation in Cebrian *et al.* including annotations of tip orientation (orange) and order parameter (*S*). (**D**) Similar data for wild-type and Spry1$^{-/-}$ kidneys. Images are from the indicated papers; model results are both based on measurements made from kidneys reported in Lefevre *et al*.

**Table S1. Kidney explant size changes following physical perturbations or tissue clearing.** Δarea (%) reflects the area change between the initial and final images, measured at 4X by brightfield microscopy (physical perturbations) or fluorescence using the E-cadherin fluorescence channel (ScaleA4/B2 tissue clearing).

| **kidney stage** | **experiment** | **n (# of samples)** | **Δarea (%, mean ± S.D.)** |
|---|---|---|---|
| E15 | control (18 hrs *ex vivo*, **Fig. 3**) | 4 | 3 ± 4% |
| E15 | + 25 nM calA (18 hrs *ex vivo*, **Fig. 3**) | 3 | 6 ± 6% |
| E15 | - ureter (18 hrs *ex vivo*, **Fig. 3**) | 6 | -1 ± 2% |
| E15 | - ureter + 25 nM calA (18 hours *ex vivo*, **Fig. 3**) | 4 | 8 ± 5% |
| E15 | cap cutting (24 hours *ex vivo* **Fig. 4**) | 6 (cap explants) | -10 ± 7% |
| E15 | cap cutting (24 hours *ex vivo* **Fig. 4**) | 3 (intact kidneys) | -2 ± 2% |
| E17 | ScaleA4/B2 tissue clearing (**Fig. 1**) | 6 | 33 ± 6% |

**Movie S1. Computational model parameters.** (**A**) Radial height occupied by tubule families in *z.* (**B**) Repulsion distance between nodes. (**C**) Planar confinement (width and depth) in the *xy* plane of tubule families each sharing a great-grandparent node.

**Movie S2. Computational model dynamics in real-time.** View of model volume from the top (*xy*), showing form-finding of tubule family spatial geometry by energy minimization.

**Movie S3. Kidney response to dispase treatment over 40 min.** (**A**) PNA lectin-labeled whole E15 kidney response to incubation in dispase imaged by live confocal fluorescence microscopy, showing (**B**) ureteric tubule delamination and retraction from the side (*xz*) and (**C**) top (*xy*).

**Movie S4. Soft material model dynamics.** (**A**) 3D printed elastic tubule family sharing a great-great-grandparent node. These families were impregnated with fluorescent methacryloxyethyl thiocarbamoyl rhodamine B. (**B**) Families arranged near the surface of a polydimethylsiloxane (PDMS) silicone elastomer block to mimic the H's tubule packing phase. (**C**) Tubule dynamics upon application of compressive strain in xy imaged by rhodamine fluorescence. (**D**) Tubule dynamics upon application of elongation strains in *z* using nylon monofilament lines secured to great-great-grandparent nodes. See Fig. 4A-D for scale bars.

**Data S1. Rhino grasshopper model file.**
